# Shape and size complementarity induced formation of supramolecular protein assemblies with metal-oxo clusters

**DOI:** 10.1101/2020.11.18.388215

**Authors:** Laurens Vandebroek, Hiroki Noguchi, Kenichi Kamata, Jeremy R. H. Tame, Luc Van Meervelt, Tatjana N. Parac-Vogt, Arnout R. D. Voet

**Affiliations:** Laboratory of Biomolecular Modelling and Design (KU Leuven), Celestijnenlaan 200G box 2403 - 3001 Leuven, Belgium; Biomolecular Architecture (KU Leuven), Celestijnenlaan 200F box 2404 - 3001 Leuven, Belgium; Laboratory of Bioinorganic Chemistry (KU Leuven), Celestijnenlaan 200F box 2404 - 3001 Leuven, Belgium; Protein Design Laboratory (Yokohama City University), Suehiro 1-7-29 - Tsurumi-ku, Yokohama 230-0045, Japan

**Author notes:** **Corresponding Author** Prof. Arnout R.D. Voet, Laboratory of Biomolecular Modelling and Design (KU Leuven), Celestijnenlaan 200G box 2403 - 3001 Leuven, Belgium.

**Keywords:** Protein Design, Polyoxometalate, crystallography, Supramolecular assembly

## Abstract

The controlled formation of protein supramolecular assemblies is challenging but it could provide an important route for the development of hybrid biomaterials. In this work, we demonstrate formation of well-defined complexes formed between the 8-fold symmetrical designer protein Tako8 and soluble metal-oxo clusters from the family of Anderson-Evans, Keggin and Zr^IV^- substituted Wells-Dawson polyoxometalates. A combination of x-ray crystallography and solution studies showed that metal-oxo clusters are able to serve as linker nodes for the bottom-up creation of protein based supramolecular assemblies. Our findings indicate that clusters with larger size and negative charge are capable of modulating the crystal packing of the protein, highlighting the need for a size and shape complementarity with the protein node for optimal alteration of the crystalline self-assembly.

## Introduction

The biocompatibility and versatility of proteins makes them ideal building blocks for the bottom-up fabrication of hybrid biomaterials with applications ranging from catalysis to drug delivery and bio-electronics.^1–3^ Supramolecular chemistry involving proteins however is a challenging area as their heterogeneous macromolecular surface makes control of host-guest interactions difficult. Nevertheless, in recent years a number of successful strategies have been developed to create symmetric protein frameworks. The group of Tezcan engineered metal binding sites at the symmetry axes of the natural protein ferritin, which serve as nodes for connection with organic linkers, reminiscent of metal organic frameworks.^4–6^ Other approaches for creating protein frameworks include the use of dimeric variants of small natural ligands to link symmetric oligomeric proteins,^7^ or macrocycles which bind to pockets on the protein surface and act as a molecular glue.^8–12^ In comparison to metal ions or organic ligands, the introduction of all-inorganic metal-oxide clusters is even more challenging because they represent non-natural molecules without evolved bio-compatibility. Polyoxometalates (POMs) are a diverse class of soluble and symmetric inorganic clusters with potential applications in various areas of chemistry and nanoscience.^13–16^ Due to their affinity towards protein surfaces, POMs have also found applications in biochemistry and medicine,^17^ and have been used as additives in protein crystallization,^18,19^ or in the development of inhibitory drug-like molecules^20^ and artificial enzymes.^21–24^

In previous work, we combined the *de novo* designed protein, a six-bladed β-propeller protein Pizza6-S with POMs with compatible symmetry in order to explore their potential for creating hybrid assemblies.^25^ The protein Pizza6-S was designed with a ring of six histidine residues around the central six-fold symmetrical cavity, creating a positively charged shallow pocket suitable for binding negatively charged polyoxometalate clusters. Dimeric Keggin POM K_11_[Ce^III^[PW_11_O_39_]_2_] was shown to bind to two Pizza6-S molecules, holding them together in solution, and altering the crystal packing of the protein into a porous framework in which the protein nodes were connected by POMs. These results demonstrated that hybrid macromolecular assemblies can be created out of symmetric proteins linked together by metal-oxo clusters, however since Pizza6-S has only a single POM binding site, formation of large dual-component supramolecular assemblies which extend further than two proteins linked by a POM ligand, is not feasible.

In this work, we set out to investigate another designer protein, Tako8, which we recently designed as a toroidal 8-fold symmetric protein with a highly polarized surface.^26^ Unlike Pizza protein, which has an axial positively-charged POM binding cavity, the surface of Tako8 was designed to be negatively-charged around the central channel, and therefore unlikely to bind POM at this position. However, the positively-charged outer rim and the 8-fold symmetry of the protein should allow for the binding of multiple POMs in a symmetric way around the torus. We demonstrate in this work that such localization of positive and the negative charges on the protein surface can result in the controlled formation of larger assemblies in which proteins are symmetrically linked via the POMs bound at the proteins periphery.

## Experimental section

### Expression and purification of Tako8 protein

A synthetic linear DNA fragment encoding Tako8 was purchased from GeneArt (Hong Kong, China), and cloned into pET28b vector (Novagen) using the NdeI and XhoI restriction sites. The protein was expressed and purified as previously described.^26^ The purified protein was concentrated to 10-20 mg/mL using a Vivaspin15R (Sartorius), and purity was estimated to be at least 95% by SDS-PAGE.

### Polyoxometalate synthesis

STA was purchased from Sigma Aldrich in the highest possible purity grade and was used without further purification. TEW^27^ and Zr-WD^28^ were synthetized and characterized according to published procedures.

### Crystallization, structure determination and refinement

Drops of 10 mg/mL Tako8 protein in 20 mM HEPES pH 8.0 and 100 mM NaCl were used in the vapor diffusion method, by mixing it in a 1:1 ratio with a precipitant solution from commercial screening kits. The conditions used for screening were chosen to be based on the native Tako8 crystallization conditions. The co-crystallization drops were identical to a native Tako8 crystallization drop, except each individual POM was added from a stock solution prepared in milliQ water. TEW was added in a concentration of 0.9 mM, STA as a concentration of 3.5 mM and a concentration of 1 mM for Zr-WD. All crystals were cryo-cooled using 20-30% glycerol as cryo-protectant. X-ray diffraction data were collected on beamline I04 of the Diamond Light Source.

Diffraction images were processed with XDS^29,30^ and scaling was performed with AIMLESS^31^. Molecular replacement using PHASER^32^ with the Pizza structure (PDB: 3WW9)^33^ as a template gave suitable solutions. Refinement was performed with PHENIX.REFINE^34,35^ and COOT^36^. The completed structures were validated with MolProbity.^37^ Data collection and refinement statistics are given in Supplementary Table 1. The coordinates and structure factor data have been deposited with the Protein Data Bank with entry codes: 6Y7N (Tako8 and TEW); 6Y7O (Tako8 and STA); 6Y7P (Tako8 and Zr-WD).

Figures were generated in PyMOL.^38^ Secondary structures were assigned with DSSP^39^ and electronic potentials were calculated using APBS and PDB2PQR^40–42^. Void volumes and percentages in the crystal packing were calculated using Mercury CSD 4.1.3 (Build 249162), using a probe of 1.2 Å and the solvent accessible surface.^43^

### Isothermal titration calorimetry

ITC experiments were carried out with a MicroCal ITC200 (Malvern). Purified Tako8 protein was placed in the cell and maintained at a temperature of 298.15K. The POM solutions were dissolved in the same buffer as the protein were injected to the cell. The buffer that was used was the same as the crystallization condition, namely 100 mM acetate pH 4.5. Instead of 3.1 M NaCl a concentration of 100 mM was used to prevent precipitation/crystallization during the experiment, while still preserving some ionic strength in solution. 18 injections of the ligand solution, 0.4 μL each, were made in total, allowing the baseline to stabilize between injections. All experiments have been performed in duplicate. The raw data were analyzed using NITPIC^44–46^, SEDPHAT^45- 47^ software with single binding site model. The figures were made with GUSSI.^48^

### Analytical Ultracentrifugation

Sedimentation velocity experiments were carried out using an Optima XL-I analytical ultracentrifuge (Beckman-Coulter, Fullerton, CA) using an An-50 Ti rotor. For sedimentation velocity experiments, cells with a standard Epon two channel centerpiece and sapphire windows were used. 400 μL of protein (38.9 μM) or 400 μL mixture solution with POMs (10, 25 and 100 μM) and 420 μL of reference buffer were used in each experiment. The same pH buffers were used as with the ITC measurement, with the addition of 100 mM NaCl in each experiment. The rotor temperature was equilibrated at 20°C in the vacuum chamber for 1.5 hours prior to starting each measurement. Absorbance (280 nm) and interference scans were collected at 10 min. intervals during sedimentation at 50,000 rpm, unless stated otherwise. The resulting scans were analyzed using the continuous distribution c(s) analysis module in the program SEDFIT.^49^ The partial specific volume of the proteins, solvent density, and solvent viscosity were calculated using the program SEDNTERP.^50^

## Results and discussion

The POM ligands used in this work belong to the three most common POM archetypes: Anderson-Evans Na_6_[TeW_6_O_24_] (TEW), Keggin H4[SiW12O40] (STA) and Zr^IV^-substituted Wells-Dawson K_15_H[Zr(α_2_-P_2_W_17_O_61_)_2_] (Zr-WD), which all exhibit multiple internal symmetry elements (Figure 1). TEW exhibits a 3-fold rotation axis, while STA has tetrahedral point symmetry, combining both 3-fold rotational symmetry with 4-fold rotation inversion symmetry. Zr-WD on the other hand can be considered as a merger of two Keggin POMs, producing a larger ellipsoid metal-oxo cluster with a 3-fold rotation axis, yet lacking any 4-fold symmetry. Since the parent Wells-Dawson POM is stable only in very acidic pH conditions, Zr^IV^-substituted Wells-Dawson was used due to its stability in a wide range of pH conditions.^51^ Moreover, it has nearly double the size of the parent Wells-Dawson POM since it consists of two Wells-Dawson units linked via a central Zr(IV) ion.

**Figure 1.**
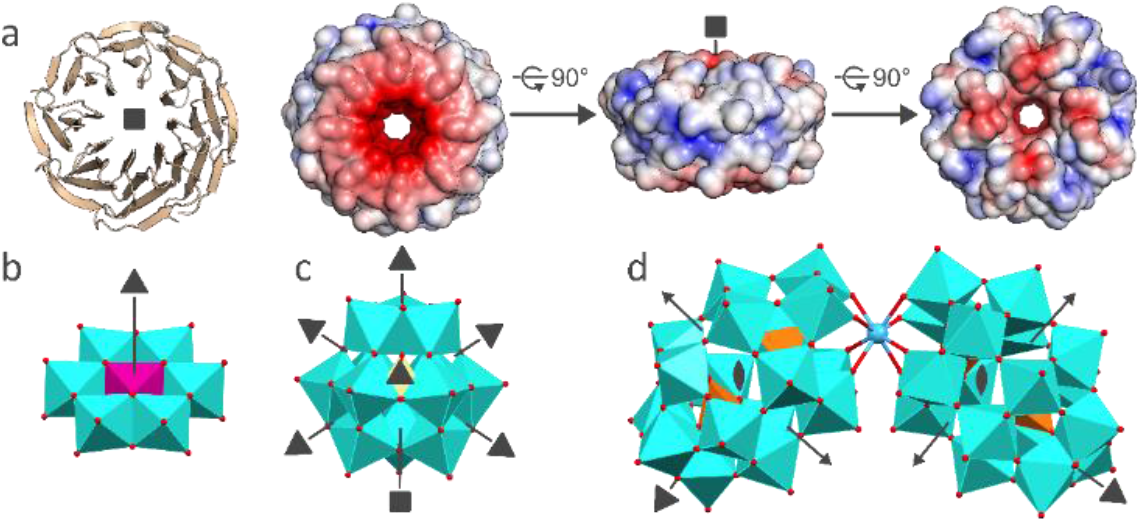
(a) Structure of Tako8 and POMs.Tako8 is a monomeric 8-fold symmetric beta-propeller with a negatively charged center while the outer ring is positively charged (cartoon representation and electrostatic surface representation of the top, side and bottom of the protein (left to right). While the backbone is designed to have 8 identical repeating blades, this translates into 4-fold symmetry in the crystal structure due to minor conformational changes of the backbone fold. Polyhedral representation of (b) the Anderson-Evans TEW Na_6_[TeW_6_O_24_], (c) Keggin STA H_4_[SiW_12_O_40_] and (d) Zr^IV^-substituted Wells-Dawson Zr-WD K_15_H[Zr(α_2_-P_2_W_17_O_61_)_2_]. Symmetry elements belonging to the protein and POMs are indicated, only for Zr-WD this symmetry reflects local symmetry and not the point group symmetry.

Tako8 alone was crystallized by the sitting drop vapor diffusion method. Crystals grown in citrate buffer at pH 5.0 and 3.4 M NaCl resulted in cubic packing of the protein. A similar arrangement was observed when acetate buffer (pH 4.5, 0.1 M) was used instead of citrate buffer. In all these forms, two Tako8 molecules are observed to pack as a face-to-face dimer. Initial attempts to crystallize Tako8/POM assemblies explored similar conditions to those used with the protein alone. STA and Zr-WD yielded crystals in the presence of acetate buffer (Supplementary Figure 1b,c). As no crystals were observed in the presence of TEW, additional screening was performed, resulting in a novel crystal form not previously observed for Tako8 alone(Supplementary Figure 1a). Unlike the cubic assembly with rhomboid voids, the new packing is built from dimers of Tako8 that stack into lamellar layers. Interestingly, while TEW induced a change in crystal packing (Supplementary Figure 2c), it could not be observed in the final crystal structure, and no evidence for its presence could be found in the electron density map or from anomalous scattering.

Both STA and Zr-WD were found in crystals grown in the presence of acetate buffer, using the anomalous signal of tungsten (Supplementary Figure 2 for STA and 3 for Zr-WD). Remarkably, a completely different packing was observed depending on the type of POM. STA was observed to bind Tako8 at two different binding sites near the outer positively charged ring (Figure 2d, 3a). The space-group and the cube-like arrangement of the protein were not altered, but STA was found to bind on the cube edges and vertices. Sitting at the center of the cube edge, STA connects two adjacent Tako8 monomers, while aligning its own 2-fold symmetry axis with the crystallographic 2-fold axis (Figure 3b).

**Figure 2.**
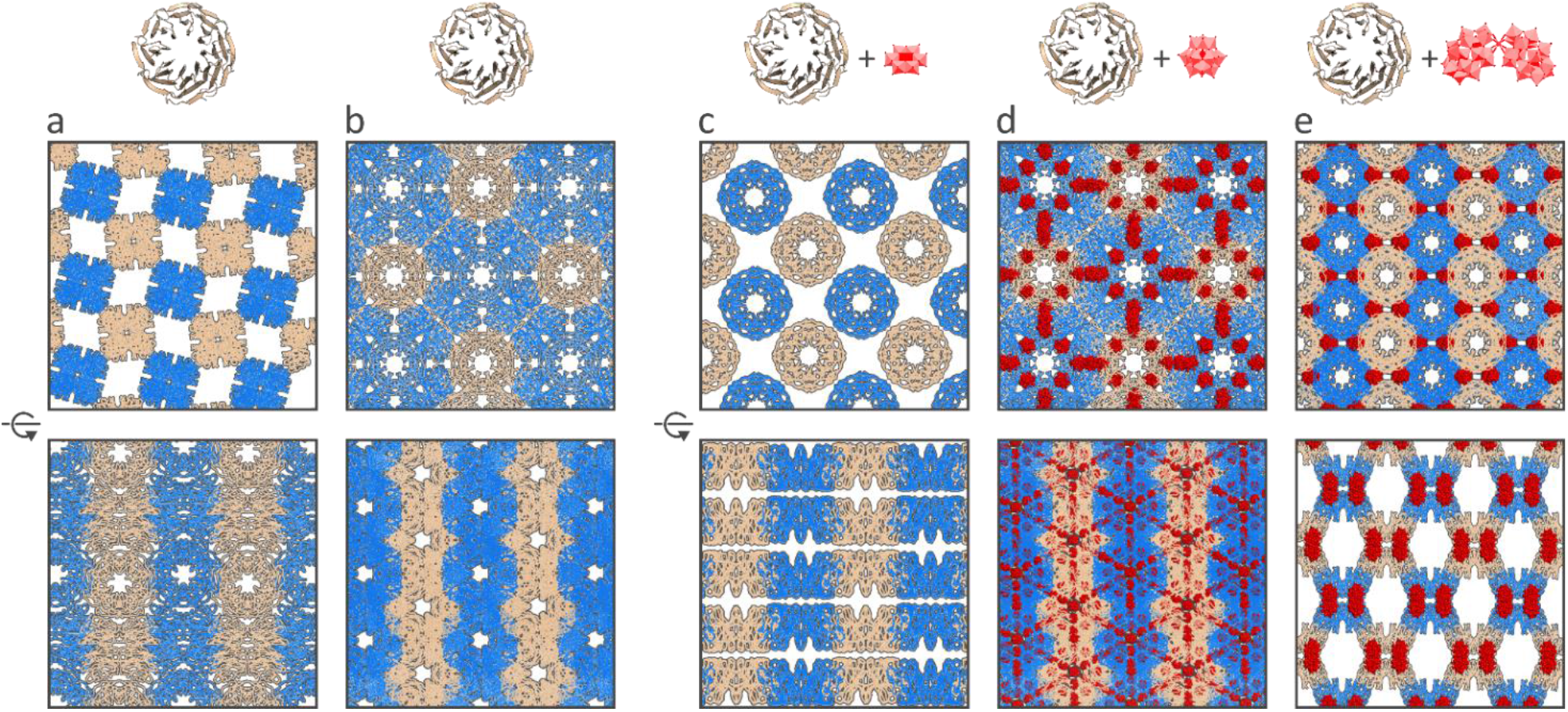
Single crystal x-ray structures of Tako8 and Tako8/POM assemblies. Proteins are represented as blue or beige ribbons to identify different layers. The POMs are represented as red spheres. The native Tako8 crystallizes into two different lattices: orthorhombic (a) and cubic (b). Addition of POMs alters the packing: in the presence of TEW a laminar stacking of dimeric Tako8 is observed while TEW is absent (c). STA does not influence the packing observed in b but binds Tako8 at the positively charged patches (d). In the case of Zr-WD the crystal packing becomes a porous honeycomb-like arrangement (e).

**Figure 3.**
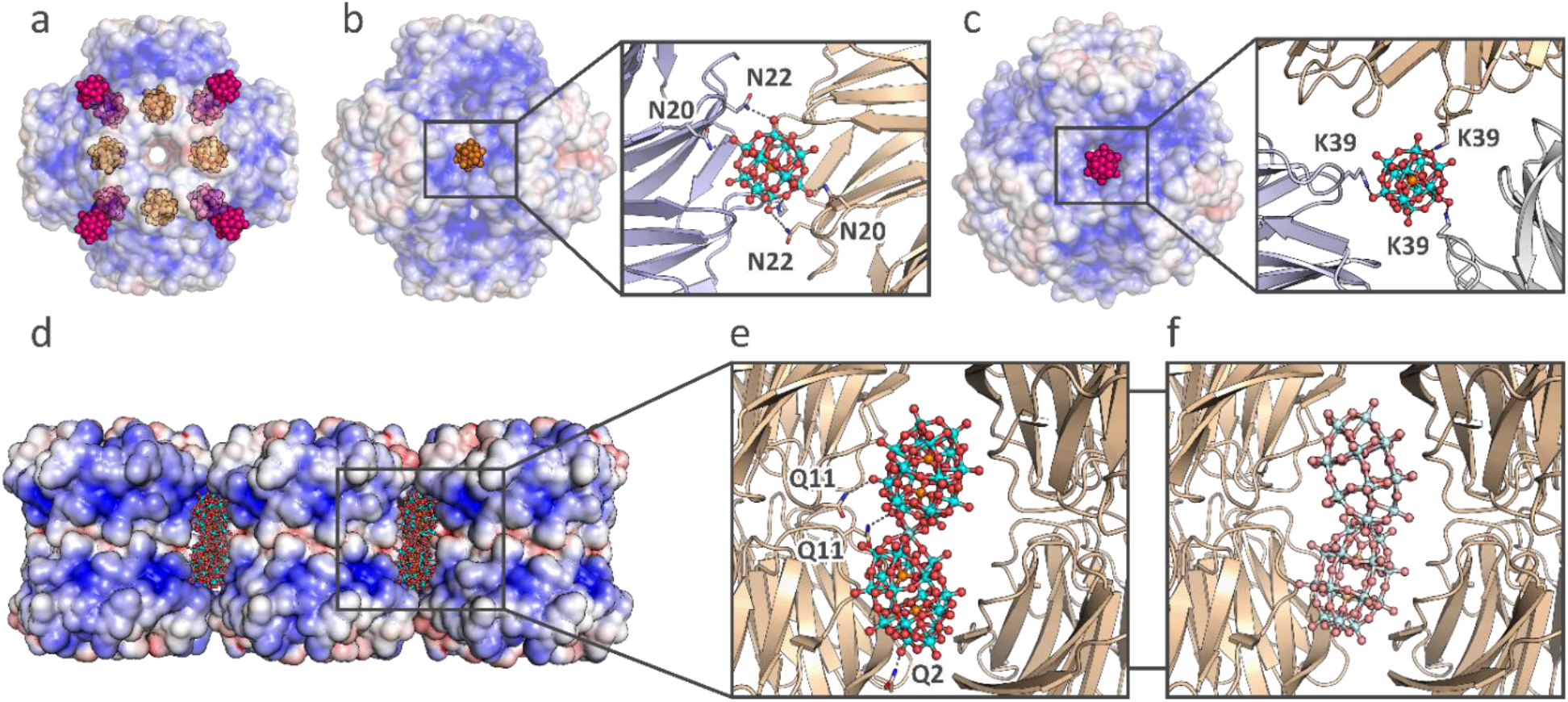
(a) Single crystal structure of Tako8 with STA. The cubic arrangement of the Tako8 in the crystal lattice is depicted with its molecular surface. The STA can be observed at two different locations: at the edges of the cube formed by two Tako8 proteins (orange spheres), and along the vertices formed by 3 Tako8 units (magenta spheres). (b) The STA at the edges binds to Asn residues and aligns with the 2-fold rotation axis of the cube. (c) The STA at the vertices is aligned along the 3-fold axis and is coordinated by 3 Lys residues (symmetry equivalents of Lys39). (d) Crystal structure of Zr-WD with Tako8. The Zr-WD (spheres) fits perfectly between two adjacent Tako8 dimers (electrostatic surface). (e) The Zr-WD is coordinated by Gln residues, and is present in two orientations at half occupancy, one more tightly bound to one set of Tako8 proteins, the other to the adjacent Tako8 dimer. (f) The alternative conformation of Zr-WD bound to the side of the Tako8 dimer, shown as a pale ball-and-stick model.

The STA interacts with the protein via hydrogen bonds with the amide group in the side chain of asparagine 20 and 22 residues (and their symmetry equivalents). This accounts for 4 out of 8 equivalent sites of a Tako8 protein being bound to POM, but STA is also observed at the other four remaining equivalent sites, which correspond to the vertices of the cube. Here, the 3-fold rotational symmetry of the POM is aligned with the 3-fold axis of the cubic assembly (Figure 3c). In this case, the STA binds to Tako8 protein via salt bridges with the positively charged lysine 39 (and its symmetry equivalents). While the final packing is porous, there are no channels linking the cubic voids, indicating that the STA and protein assemble together into the crystal, rather than STA binding subsequently to a preformed protein lattice.

Unexpectedly, and in contrast to STA, Zr-WD profoundly changed the crystal packing of the protein despite identical chemical composition of the crystallization buffer. The face-to-face dimer, common to all Tako8 crystal structures, is still present, but the packing with Zr-WD results into a highly porous honeycomb arrangement (Figure 2e). Zr-WD is bound to both Tako8 rings in the protein dimer at a positively charged surface. The only directed interactions between POM and Tako8 are between oxygen atoms of the POM and the amide side-chains of glutamine 11 and glutamine 2, and their symmetry equivalents (Figure 3d,e). In contrast to STA, which is bound on the bottom of Tako8, Zr-WD instead binds on the periphery of the Tako8 dimer and spans both monomers (Figure 4).

**Figure 4.**
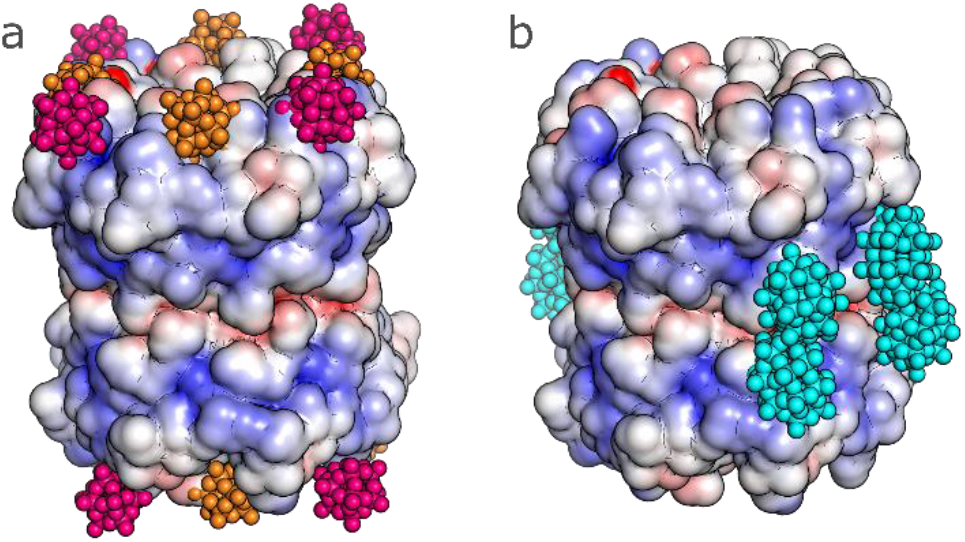
Comparison of the POM binding sites on the surface of the Tako8 dimer between STA (a) and Zr-WD (b). The orientation of the Tako8 dimer is identical in both images. The POMs are shown as colored spheres, where STA bound on the edge of the cube are colored orange, STA bound on the vertices of the cube are colored magenta and Zr-WD is colored cyan.

Interestingly, Zr-WD is commonly observed as the cis sandwich-type in small molecule crystal structures, but here it is observed in the trans-conformation.^28^ The dynamic nature of this sandwich-type POM apparently allows it to change shape to maximize its contact with the positively charged protein surface. The trans-conformation of the POM prevents it from forming the edge-to-edge interactions with Tako8 molecules seen in the cubic packing (Supplementary Figure 4). As expected, the negatively-charged central cavity proved unfavorable for binding POMs, despite its suitable shape and symmetry. POMs were found to bind solely to the positively-charged sides of Tako8, and the interaction were largely electrostatic in nature and involved very few hydrogen bonds.

The affinities of the POMs for Tako8 in solution were measured by isothermal titration calorimetry (ITC) and analytical ultracentrifugation (AUC) (Supplementary Figure 5). As expected, no binding could be observed for TEW (Supplementary Figure 6a), while STA binds Tako8 with moderate affinity (K_d_ = 57.8 μM, see Supplementary Figure 6b) compared to Zr-WD(K_d_ = 1.13 μM, see Supplementary Figure 6c). The results suggest that not only the overall charge of the POM influences the affinity of binding, but that the POM size and shape also play a crucial role. TEW is the smallest POM in the series, while STA is intermediate in size, and Zr-WD is the largest with the highest charge, thus allowing for the largest number of non-covalent interactions.^28^ Both STA and Zr-WD make few direct interactions compared to the previously reported Pizza6-S:STA complex which had stronger affinity. The strength of the interaction could thus presumably be increased by introducing more positive charge or more hydrogen bond forming residues at the POM binding sites. AUC confirmed the absence of any interaction between TEW and Tako8. On the contrary, the complexes formed with STA and Zr-WD were very large and immediately sedimented at the onset of centrifugation (Supplementary Figure 7). This is consistent with the observed crystal structures which showed that both STA and Zr-WD are able to link multiple Tako8 proteins, and implies that in solution such binding results in extensive networks with very high molecular weight.

## Conclusion

Despite the highly symmetric nature of POMs, our results demonstrate that protein-based supramolecular networks can be created by combining symmetrical proteins and soluble metal-oxo clusters with matching shape, instead of matching symmetry. The toroidal Tako8 dimerizes at mildly acidic pH and such dimer has a large positively-charged surface at its periphery, which matches in shape and size the Zr-WD oxo-cluster. The combined solution and crystallographic studies indicate that charge and shape complementarity drive the formation of Tako8-POM assemblies. The smaller and less negatively charged metal-oxo clusters do not dictate the protein crystal packing proteins and sit at positive patches in the crystalline assembly. On the other hand, larger metal-oxo clusters such as Zr-WD are able to bind with high affinity to positively-charged surface patches of proteins, and control their self-association. They act as an “electrostatic glue”, connecting the positively-charged surfaces of adjacent symmetric protein molecules in a plane perpendicular to the Tako8 protein’s symmetry axis. The dynamic nature of the Zr-WD metal-oxo cluster allows it to change conformation in order to optimize the linking of two partner proteins without even matching the POMs symmetry. Further rational design of protein binding surfaces with complementary shape and charge to metal-oxo clusters will allow the facile construction of both 2D and 3D supramolecular assemblies that combine the properties of both biological and inorganic components.

## Supporting information

Tako8 + TEW - Crystal Structure

Tako8 + STA - Crystal Structure

Tako8 + Zr-WD - Crystal Structure

## ASSOCIATED CONTENT

The following files are available free of charge.

Crystal structures of the Tako8 crystals grown in the presence of TEW (PDB ID: 6Y7N), STA (PDB ID: 6Y7O) and Zr-WD (PDB ID: 6Y7P) (CIF)

Experimental information, supplementary figures and table (PDF)

## AUTHOR INFORMATION

### Author Contributions

The manuscript was written through contributions of all authors. LV and HN purified and crystallized all complexes, LV and KK performed AUC and ITC. TPV, JRHT LVM and ARDV conceived and supervised all experiments.

## Funding Sources

This research was supported by Research Foundation Flanders (FWO) project grants (G0E4717N, G0F9316N and G051917N) and the PhD fellowship of LV.

## ACKNOWLEDGMENT

This research was supported by Research Foundation Flanders (FWO) project grants (G0E4717N, G0F9316N and G051917N) and the PhD fellowship of LV. JRHT thanks OpenEye Scientific Software for support. The authors thank the Diamond Light Source for beam time allocation, and the staff of the I-04 beamline for assistance with data collection.

## ABBREVIATIONS

POM: polyoxometalate
TEW: tellurotungstic Anderson-Evans
STA: silicotungstic acid Keggin
Zr-WD: Zr^IV^-substituted phosphotungstic Wells-Dawson
ITC: isothermal titration calorimetry
AUC: analytical ultracentrifugation.

---

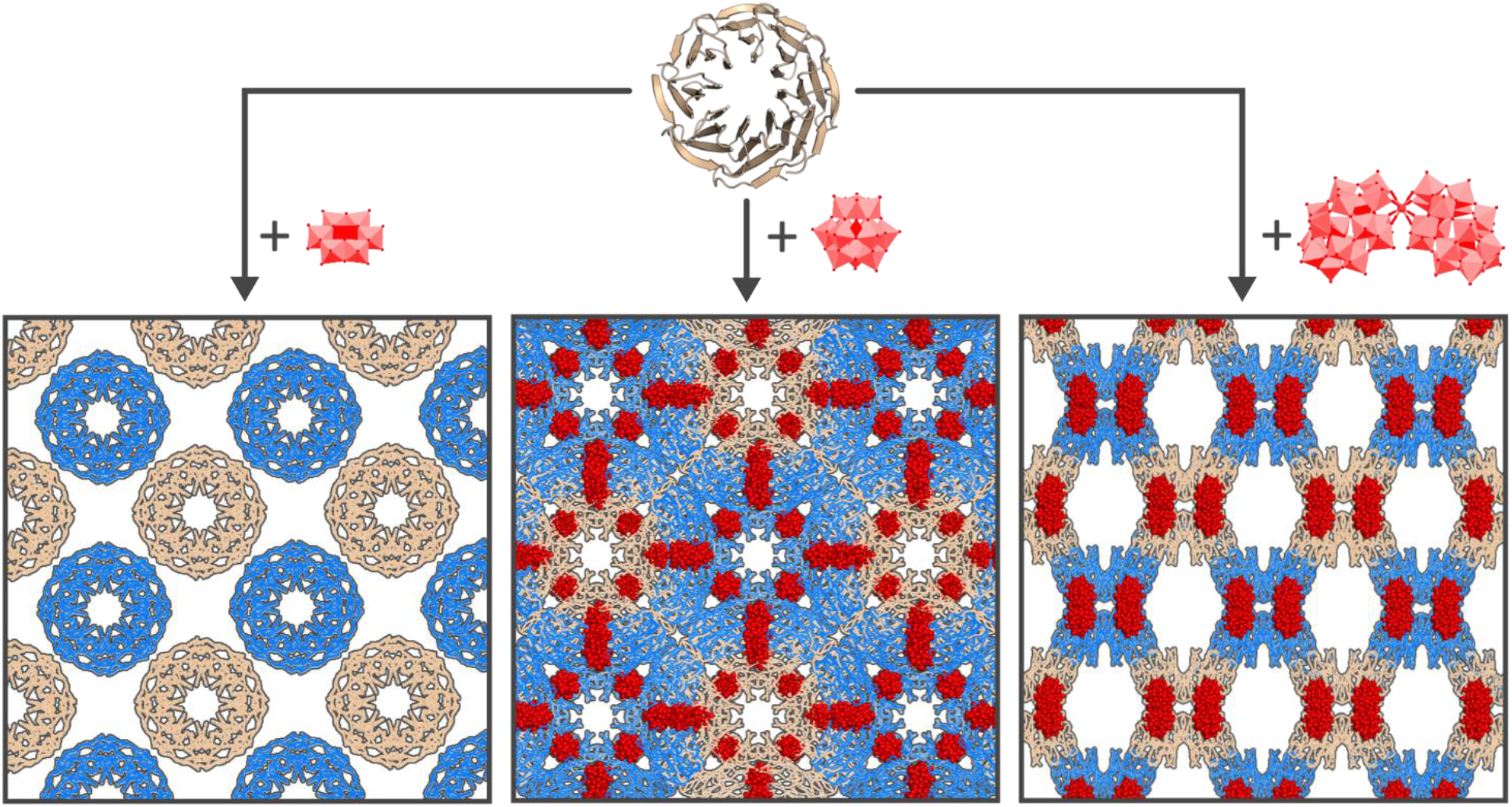
For Table of Contents Only.

## Notes

### Competing Interest Statement

The authors have declared no competing interest.

