## Supplementary Information for "Shape and size complementarity induced formation of supramolecular protein assemblies with metal-oxo clusters"

### Supplementary figures and tables

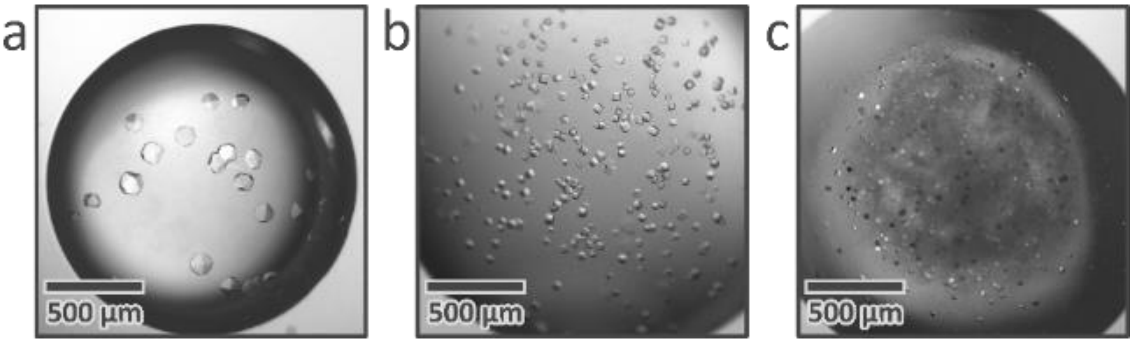

Supplementary Figure 1. Microscopic images of the Tako8 crystals obtained in the presence of TEW (a), STA (b) and Zr-WD (c).

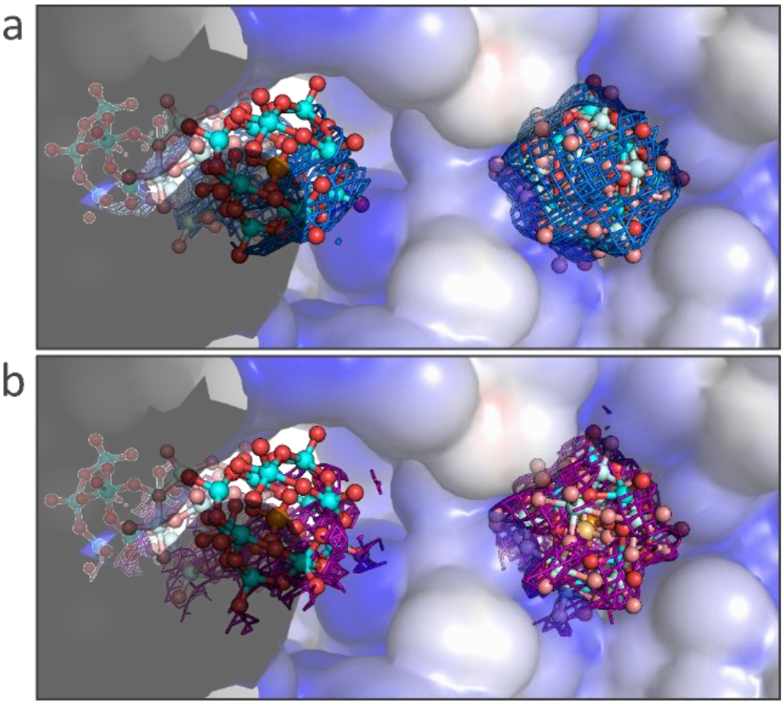

Supplementary Figure 2. $2F_o$-$F_c$ electron density map (a) and anomalous difference map (b), shown as a blue and purple mesh respectively at 2.0 σ, of the channel-bound STA (left) and ring-bound STA (right). An alternative conformation of the POM is shown as a pale-colored ball-and-stick model.

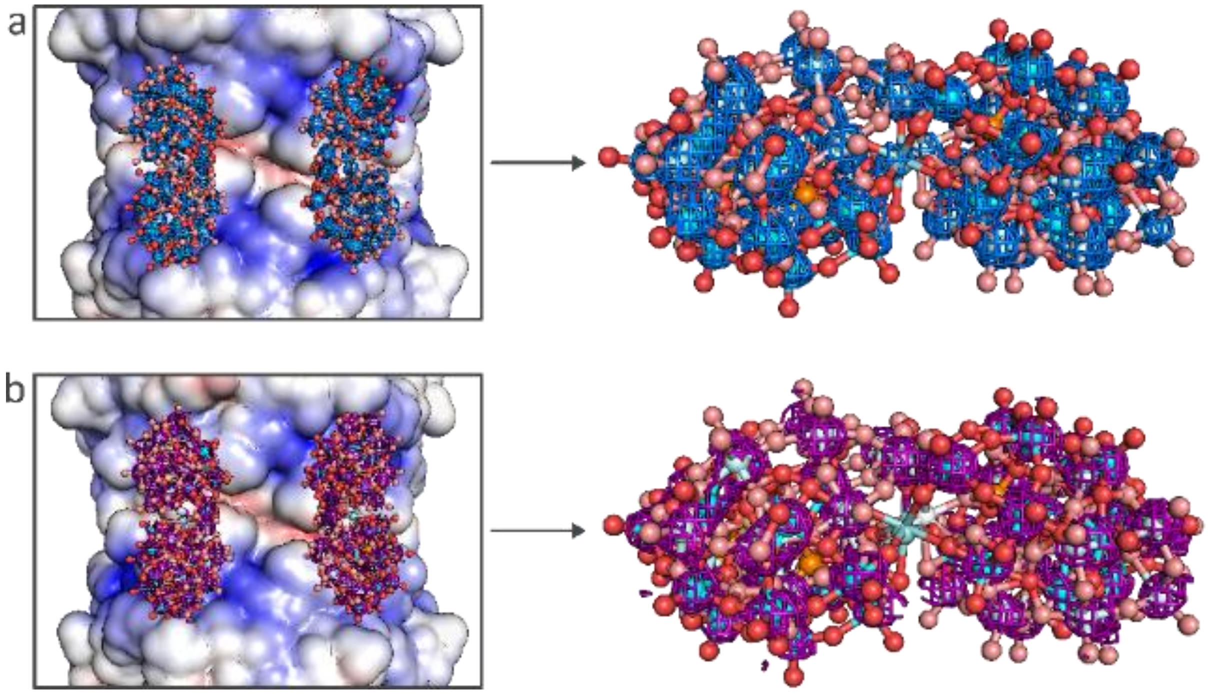

Supplementary Figure 3. $2F_o$-$F_c$ electron density map (a) and anomalous difference map (b), displayed as a blue and purple mesh respectively at 2.0 σ, show the presence of Zr-WD in the crystal structure. Zr-WD is present in two alternative orientations, each having an occupancy of 40% or 80% in total.

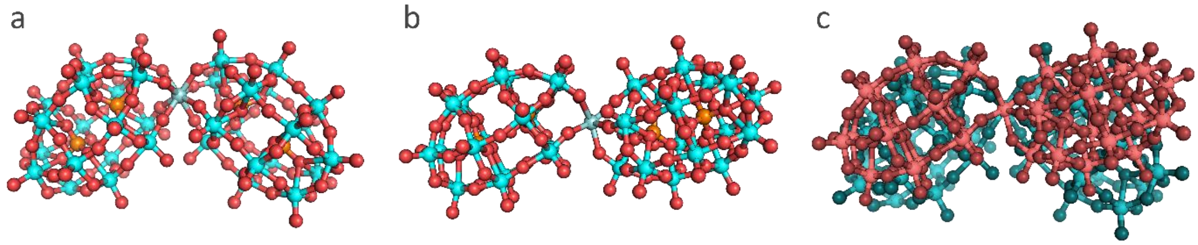

Supplementary Figure 4. Side-by-side comparison of the Zr-WD structure in the inorganic crystal structure[1] (a) and as observed in the Tako8 co-crystal (b). A manual overlay between the inorganic structure (blue) and the co-crystallized structure (red) clearly shows the difference in the orientation of the Wells-Dawson subunits (c).

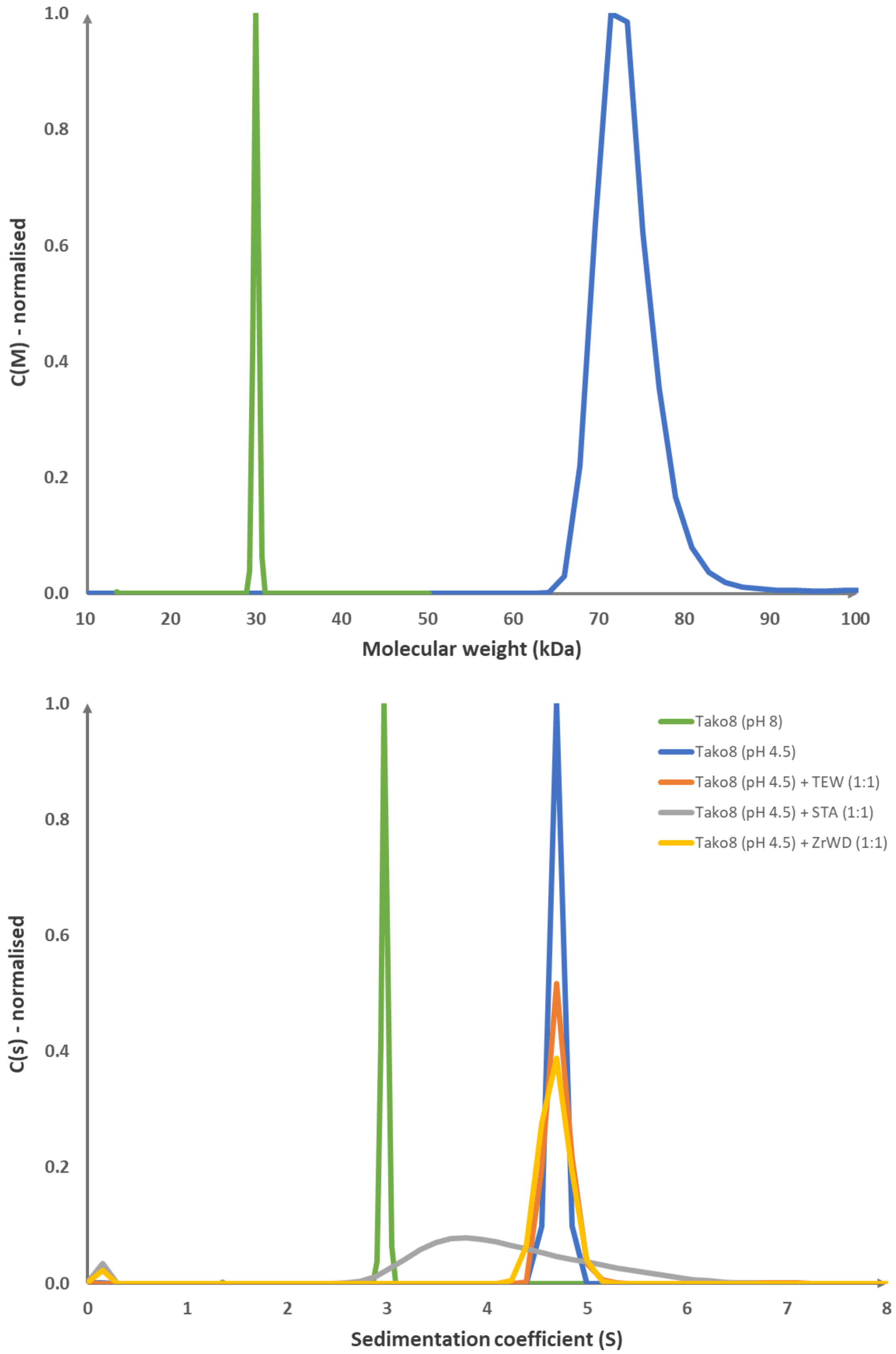

Supplementary Figure 5. Analytical ultracentrifugation spectra of Tako8 and its complexes with TEW, STA and Zr-WD. The molecular weight distribution of Tako8 at 30,000 rpm, the green line is the Tako8 monomer at pH 8 as observed in previous work and the blue line is the dimer formed at pH 8 (top); the sedimentation distribution of Tako8 and equimolar mixtures with TEW, STA and Zr-WD at 50,000 rpm (bottom).

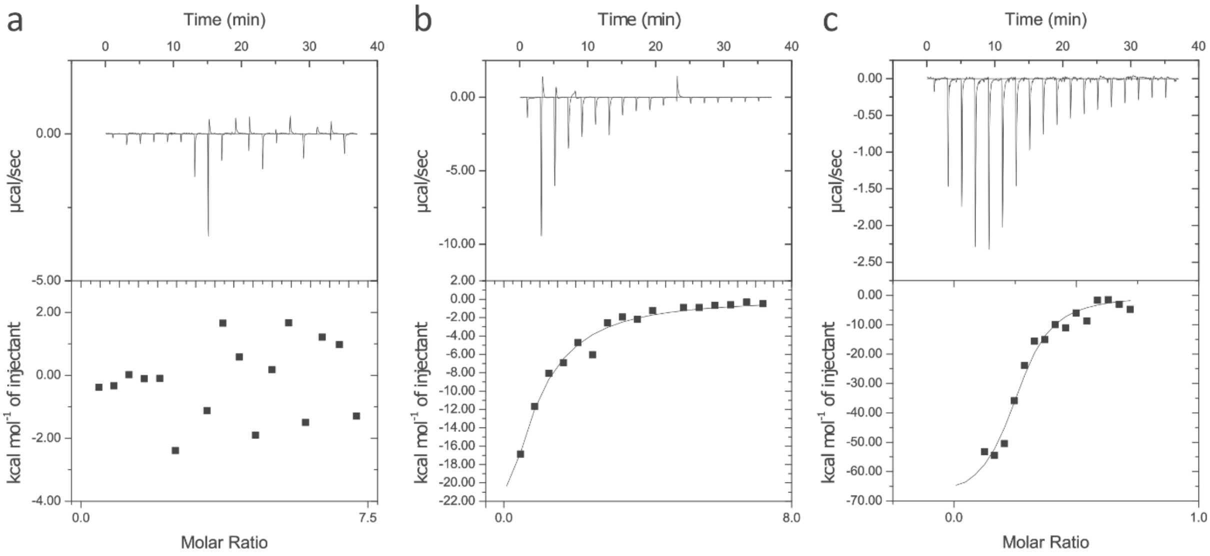

Supplementary Figure 6. ITC thermograms of POM interaction with Tako8. No interaction could be observed for TEW (a), while STA has a low affinity (b) and Zr-WD has a higher affinity (c).

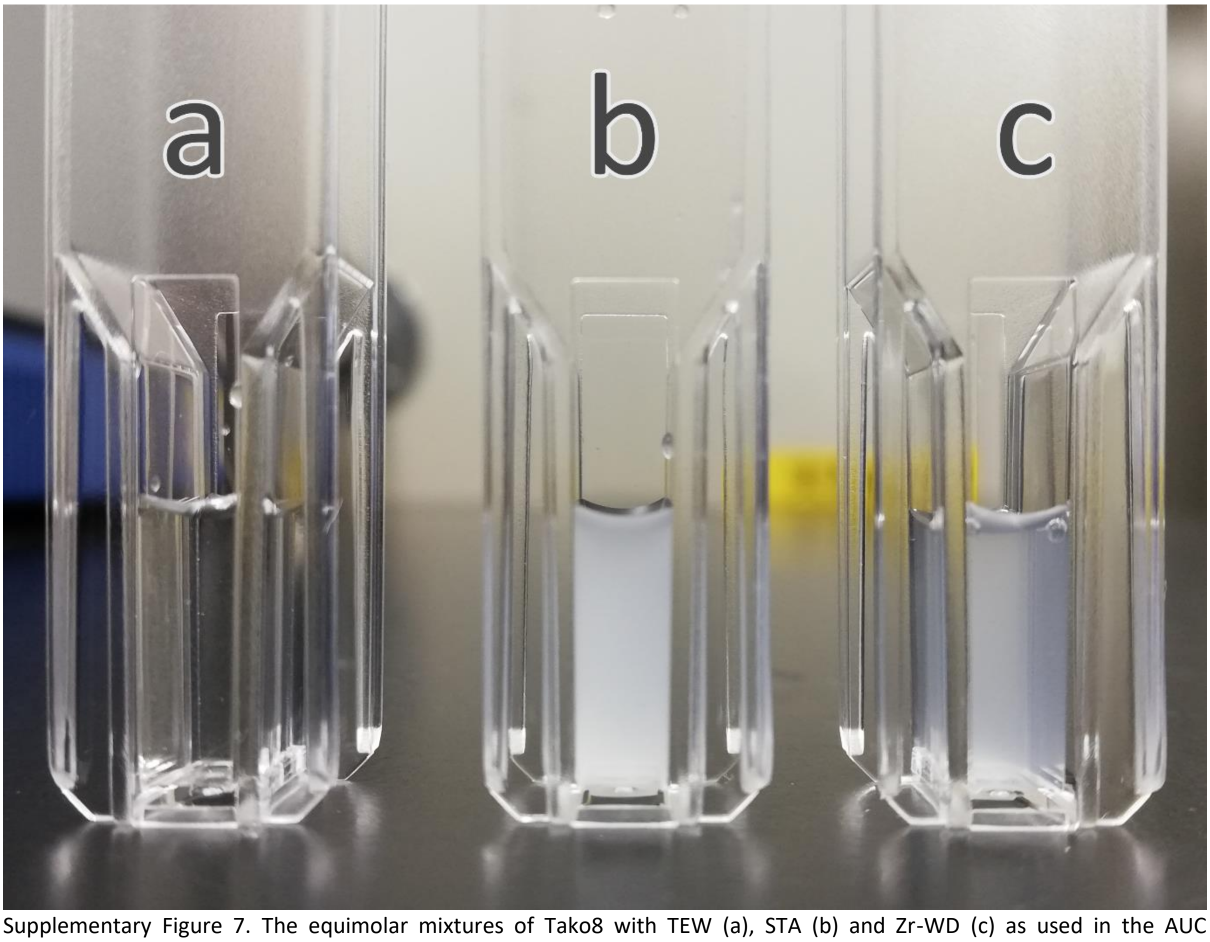

Supplementary Figure 7. The equimolar mixtures of Tako8 with TEW (a), STA (b) and Zr-WD (c) as used in the AUC measurements. Precipitation of the Tako8/STA and Tako8/Zr-WD complex is clearly visible.

Supplementary Table 1. X-ray data collection and refinement statistics.

| | Tako8 + TEW* | Tako8 + STA | Tako8 + Zr-WD |
|---|---|---|---|
| **PDB code** | **6Y7N** | **6Y7O** | **6Y7P** |
| **Data collection** | | | |
| Diffraction source | DLS, I-04 | DLS, I-04 | DLS, I-04 |
| Wavelength (Å) | 0.9795 | 0.9795 | 0.9795 |
| Resolution range (Å) | 46.93-1.60 (1.63-1.60) | 47.28-2.30 (2.38-2.30) | 46.83-1.75 (1.81-1.75) |
| Space group | *P2₁* | *C2* | *C2* |
| *a*, *b*, *c* (Å) | 73.59, 60.93, 73.61 | 94.55, 133.50, 81.75 | 107.35, 72.8, 54.29 |
| α, β, γ (°) | 90.00, 90.00, 90.00 | 90.00, 125.27, 90.00 | 90.00, 120.38, 90.00 |
| Reflections (measured/unique) | 283160/85806 | 359925/36727 | 454003/70938 |
| Completeness (%) | 99.5 (99.6) | 99.9 (100) | 99.2 (99.8) |
| Mean $I/\sigma(I)$ | 21.4 (8.3) | 8.3 (2.5) | 10.8 (2.5) |
| Multiplicity | 3.3 | 9.8 | 6.4 |
| $R_{pim}$ ‡ | 0.023 (0.070) | 0.090 (0.679) | 0.0587 (0.459) |
| CC(1/2)§ | 0.997 (0.994) | 0.989 (0.762) | 0.996 (0.810) |
| Wilson B factor (Å$^2$) | 14.1 | 36.8 | 17.0 |
| **Refinement statistics** | | | |
| Resolution range (Å) | 46.93-1.60 | 47.28-2.30 | 46.83-1.75 |
| *R* factor$^{\Pi}$/$R_{free}$$^{\Pi}$ | 0.217/0.250 | 0.241/0.293 | 0.178/0.215 |
| No. of atoms in structure | | | |
| Protein | 4874 | 7222 | 2449 |
| Ligand | 0 | 848 | 324 |
| Water | 596 | 65 | 313 |
| R.m.s. deviations from ideal | | | |
| Bond lengths (Å) | 0.003 | 0.012 | 0.008 |
| Bond angles (°) | 0.68 | 1.198 | 0.880 |
| Ramachandran plot, residues in (%) | | | |
| Most favorable region | 93.22 | 91.07 | 92.11 |
| Additional allowed region | 6.78 | 8.61 | 7.89 |
| Average *B* factor (Å$^2$) | | | |
| Main chain (A) | 15.5 | 36.9 | 17.5 |
| Side chains (A) | 21.8 | 39.9 | 23.4 |
| Main chain (B) | 15.6 | 37.2 | / |
| Side chains (B) | 21.7 | 40.3 | / |
| Main chain (C) | / | 37.0 | / |
| Side chains (C) | / | 39.8 | / |
| POMs | / | 19.3 | 34.9 |
| Waters | 28.2 | 53.5 | 24.8 |

*: The TEW POM was not observed in the crystal structure. ‡: $R_{pim} = \sum_{hkl} [1/(N-1)]^{1/2} \sum_i |I_i(hkl) - \langle I(hkl)\rangle| / \sum_{hkl} \sum_i I_i(hkl)$, where $I_i(hkl)$ is the intensity of an observation, $\langle I(hkl)\rangle$ is the mean value for that reflection and the summations are over all reflections. §: $CC(1/2) = \sum (x-\langle x\rangle)(y-\langle y\rangle)/[\sum (x-\langle x\rangle)^2 \sum(y-\langle y\rangle)^2]^{1/2}$ Π: R-factor = $\sum_{hkl} ||F_{obs}| - |F_{calc}|| / \sum_{hkl} |F_{obs}|$, where $F_{obs}$ and $F_{calc}$ are the observed and calculated structure-factor amplitudes, respectively. The free R-factor was calculated with 5% of the data excluded from the refinement. Values in brackets are for the outer shell.

Supplementary Table 2. Matthews coefficient and calculated volume and percentage of voids in the crystal packings

| Crystal | Matthews Coefficient | Calculated voids in the crystal packing | |
|---|---|---|---|
| | | Percentage within unit cell | Volume ($nm^3$) |
| Tako8 (*P*42 21 2)* | 2.68 | 50.0% | 375.1 |
| Tako8 (*C*2)§ | 1.95 | 38.7% | 316.4 |
| Tako8 + TEW | 2.33 | 41.5% | 137.0 |
| Tako8 + STA | 1.98 | 38.5% | 324.1 |
| Tako8 + Zr-WD | 2.58 | 43.4% | 158.9 |

*PDB ID: 6G6M

§PDB ID: 6G6N
